# Plant defense resistance in natural enemies of a specialist insect herbivore

**DOI:** 10.1101/710681

**Authors:** Xi Zhang, Cong van Doan, Carla C.M. Arce, Lingfei Hu, Sandra Gruenig, Christian Parisod, Bruce E. Hibbard, Maxime Hervé, Christelle A.M. Robert, Ricardo A.R. Machado, Matthias Erb

## Abstract

Plants defend themselves against herbivores through the production of toxic and deterrent metabolites. Adapted herbivores can tolerate and sequester these metabolites, allowing them to feed on defended plants and become toxic to their own enemies. Can herbivore natural enemies overcome sequestered plant defense metabolites to prey on adapted herbivores? To address this question, we studied how entomopathogenic nematodes cope with benzoxazinoid defense metabolites that are produced by grasses and sequestered by a specialist maize herbivore, the western corn rootworm. We find that nematodes from US maize fields in regions in which the western corn rootworm was present over the last 50 years are behaviorally and metabolically resistant to sequestered benzoxazinoids and more infective towards the western corn rootworm than nematodes from other parts of the world. Exposure of a benzoxazinoid-susceptible nematode strain to the western corn rootworm for five generations results in higher behavioral and metabolic resistance and benzoxazinoid-dependent infectivity towards the western corn rootworm. Thus, herbivores that are exposed to a plant defense sequestering herbivore can evolve both behavioral and metabolic resistance to plant defense metabolites, and these traits are associated with higher infectivity towards a defense sequestering herbivore. We conclude that plant defense metabolites that are transferred through adapted herbivores may result in the evolution of resistance in herbivore natural enemies. Our study also identifies plant defense resistance as a novel target for the improvement of biological control agents.

## Introduction

Despite the high abundance and diversity of arthropod herbivores, plants dominate terrestrial (agro)-ecosystems (1). Predation by herbivore natural enemies and plant defenses are thought to contribute to this phenomenon (2, 3). Plants defend themselves against herbivores using a variety of strategies, including the production of specialized defense metabolites that are toxic and/or reduce their attractivity and digestibility (4–7). However, many herbivores evolved mechanisms to overcome the negative effects of plant defense metabolites, including behavioral avoidance, excretion, target site insensitivity and detoxification through conjugation and breakdown (6, 8, 9). As a result, herbivores are often able to feed on defended plants and to ingest plant toxins without suffering major fitness consequences.

The ability to tolerate plant toxins has also enabled some specialized herbivores to co-opt plant defense metabolites for self-defense against their own natural enemies (10, 11). Sequestration of plant toxins as a form of adaptation is relatively widespread in specialized insect herbivores (12, 13). Plant toxins may also accumulate in non-adapted insect herbivores, which are often inefficient at metabolizing and/or detoxifying plant defense compounds (14–16). Consequently, predators, parasites and parasitoids are often exposed to plant toxins as they feed on herbivores. Despite the fact that plant toxin exposure of the third trophic level is common in nature, herbivore natural enemies succeed at controlling herbivores and reduce their negative impact on plant fitness and yield (3). How top-down control of herbivores is maintained in the face of the abundance, diversity and ubiquity of plant defense metabolites is a potentially important open question in multitrophic interaction research and chemical ecology.

One possible explanation for the success of herbivore natural enemies is that, similar to herbivores, they may have evolved the capacity to resist or tolerate plant defense metabolites (17). Different degrees of resistance to plant toxins have been observed in predators and parasitoids (18–21). However, whether plant defense metabolites can drive the evolution of resistance of members of the third trophic level, and to what extent resistance to plant defense metabolites improves the capacity of herbivore natural enemies to prey on adapted herbivores, is not well understood.

To address the questions above, we studied the impact of plant-derived benzoxazinoids on entomopathogenic nematodes. Benzoxazinoids are multifunctional defense metabolites that are produced by grasses such as wheat and maize (22) and protect them against generalist herbivores (23–25). The western corn rootworm (*Diabrotica virgifera virgifera*), a specialized maize herbivore and important agricultural pest, is fully resistant to benzoxazinoids (26). The rootworm larvae are attracted to benzoxazinoids (27) and accumulate them in their bodies (28). Entomopathogenic nematodes such as *Heterorhabditis bacteriophora* are common in natural and agricultural ecosystems across the globe and co-occur with the western corn rootworm in some areas. They are used as biological control agents against many different root pests, including the western corn rootworm (29, 30). Benzoxazinoid sequestration by the western corn rootworm reduces the capacity of a commercial *H. bacteriophora* strain to infect and kill the herbivore, suggesting that the corn rootworm co-opts these plant defense metabolites for self-protection against entomopathogenic nematodes (28). Using natural variation and forward evolution, we investigated whether adapted *H. bacteriophora* nematodes are able to overcome this defenses strategy of the western corn rootworm, and whether their capacity to resist benzoxazinoids is associated with increased infectivity towards the western corn rootworm.

## Results

To test whether entomopathogenic nematodes that share an evolutionary history with the western corn rootworm may be able to resist benzoxazinoids, we established a global collection of 25 *Heterorhabditis spp*. strains, including strains collected from regions in which the western corn rootworm has been present for more than 50 years (henceforth called the primary range), and strains collected from other regions of the world in which the western corn rootworm is not present or has not been present until recently (Fig. 1A, table S1). Nematodes from the primary range where isolated from maize fields in which the western corn rootworm is regularly present. They are thus likely to have encountered this herbivore in the past. *H. bacteriophora* has a broad host range, and the strains may also have infected other root herbivores occurring in maize fields, including wireworms and other rootworm species. Benzoxazinoids and their breakdown products can be found in the midgut of a wide range of insect herbivores (31, 32), but the western corn rootworm is the only herbivore known to selectively accumulate benzoxazinoids in its hemolymph (28). Nematode strains from other parts of the world never encountered the western corn rootworm, as they came from regions where the rootworm is not present, or they were isolated before the western corn rootworm invaded these regions (table S1).

**Fig. 1.**
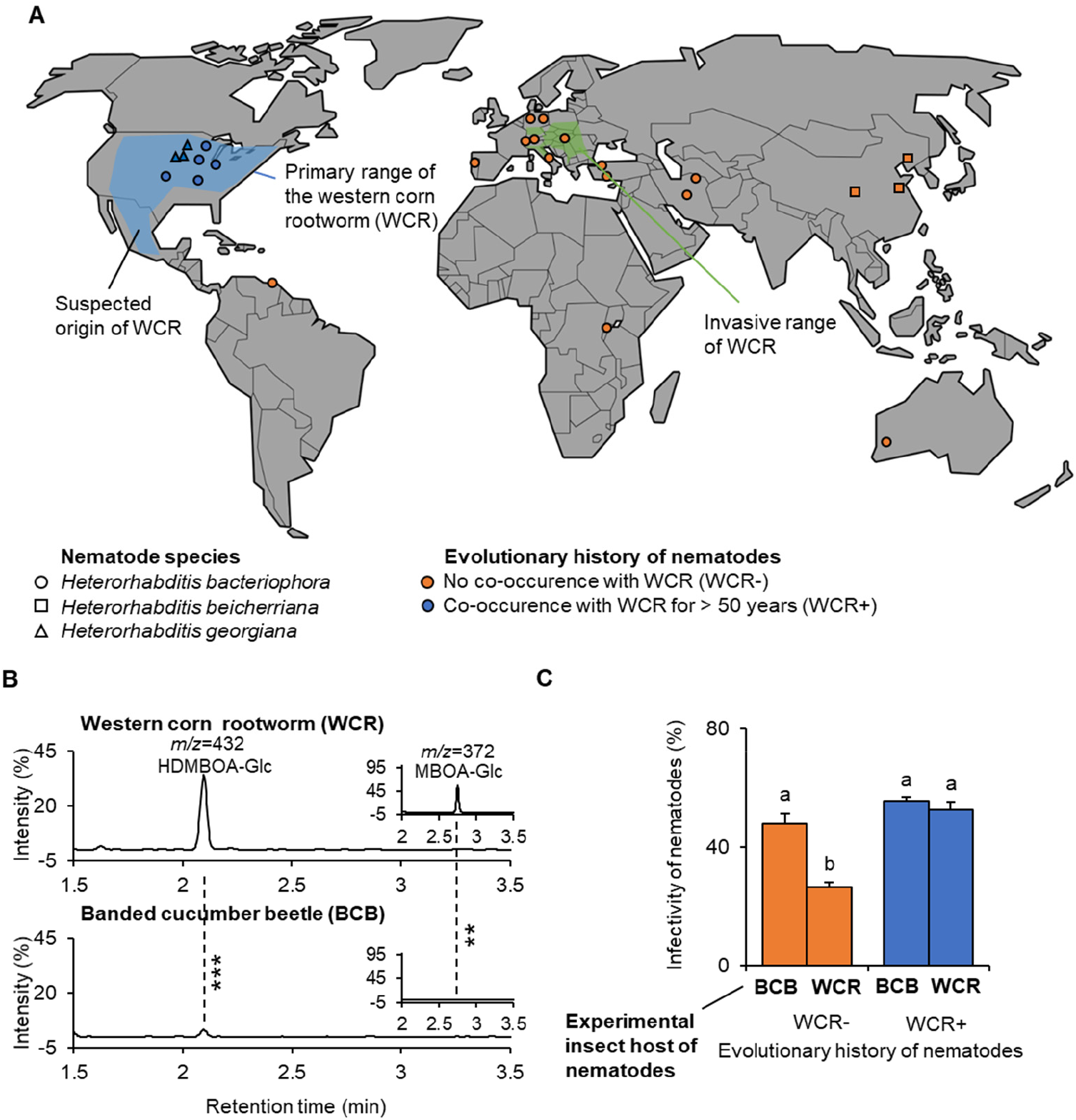
Entomopathogenic nematodes from the primary range of the benzoxazinoid-sequestering western corn rootworm are more infective towards the western corn rootworm than the non-sequestering banded cucumber beetle. (**A**) World map showing the origin of the collected entomopathogenic nematode strains together with the primary and invasive ranges of the western corn rootworm, a specialized maize herbivore (WCR; data from 2012). Note that nematode strains from the invasive range do not share any evolutionary history with WCR, as they were collected before invasion. For detailed information about the different strains, refer to table S1. (**B**) Chromatograms of plant-derived benzoxazinoids in the body of WCR (top), and the banded cucumber beetle (BCB, bottom) a generalist root herbivore which does not sequester benzoxazinoids and is mainly present in Central America, Mexico and the Southern US, outside of the nematode sampling range. Asterisks indicate significant differences between herbivore species (*P* <0.001). For quantitative comparisons, refer to Fig. S1. (**C**) Infectivity of nematodes towards WCR and BCB. Average (+SEM) infectivity is shown for nematodes with an evolutionary history with WCR of more than 50 years (blue) and nematodes without evolutionary history with WCR (orange). Different letters indicate significant differences between treatments (False discovery rate corrected *P* <0.05).

Using internal transcribed spacer rRNA gene sequencing, 19 nematode strains within our collection were confirmed to be *H. bacteriophora*. Three strains from China were re-classified as *H. beicheriana*, and three strains from the US were identified as *H. georgiana*, both of which are closely related to *H. bacteriophora*. In a first experiment, we compared the infectiveness of the different nematode strains towards larvae of the western corn rootworm and larvae of the banded cucumber beetle (*D. balteata*). In contrast to the western corn rootworm, the banded cucumber beetle feeds on many different plant species apart from maize and does not sequester benzoxazinoids (Fig. 1B and S1) (28). The banded cucumber beetle occurs mainly in Central America, Mexico and the Southern US, outside of our collection range. Thus, none of the tested nematode strains are likely to share an evolutionary history with this herbivore.

Infectivity tests revealed a significant interaction between the host herbivore species and the evolutionary history of the nematodes (Fig. 1C, table S3). Nematode strains from the primary range of the western corn rootworm were able to infect and kill the western corn rootworm and the banded cucumber equally well. By contrast, the infectivity of nematodes from outside the primary range was significantly lower for the western corn rootworm than the banded cucumber beetle (Fig. 1C). Thus, nematodes that evolved in the presence of the western corn rootworm have an increased capacity to infect and kill this specific herbivore. This pattern was similar when only strains belonging to *H. bacteriophora* were considered (Fig. S2). The infectivity of *H. georgiana* strains, all of which come from the primary range of the western corn rootworm, was higher towards the western corn rootworm than the infectivity of *H. beicheriana* strains, which come from Asia where the western corn rootworm is not present (Fig. S2).

The banded cucumber beetle and western corn rootworm differ in many traits apart from benzoxazinoids that may explain the pattern observed in Fig. 1. To specifically test for the role of benzoxazinoids in determining the higher infectivity of nematode strains from the primary range of the western corn rootworm, we exposed the different nematode strains to western corn rootworm larvae that previously fed on wild type or benzoxazinoid-deficient *bx1* maize mutant plants. *Bx1*-mutant fed western corn rootworm larvae accumulate only low amounts of benzoxazinoids (Fig. 2A and S3) (28). Nematodes from the primary range were able to infect western corn rootworm larvae equally well, independently of whether the larvae fed on benzoxazinoid-containing wild type or benzoxazinoid-deficient *bx1* mutant maize roots (Fig. 2B). By contrast, nematodes from other parts of the world suffered from a suppression of infectivity when exposed to larvae fed on wild type plants compared to *bx1* mutant fed larvae (Fig. 2B). This pattern was largely consistent across strains (Fig. S4). The benzoxazinoid susceptibility of the different nematode strains (i.e. the difference in infectivity towards wild type and *bx1*-fed western corn rootworm larvae) was negatively correlated to their infectivity towards the western corn rootworm (as measured in the previous experiment; Fig. 2), but not correlated to their infectivity towards the banded cucumber beetle (Fig. S5). Thus, nematodes from the primary range are less susceptible to the benzoxazinoid-dependent defenses of the western corn rootworm than nematodes from other parts of the world, and this trait is associated with higher infectivity towards the western corn rootworm.

**Fig. 2.**
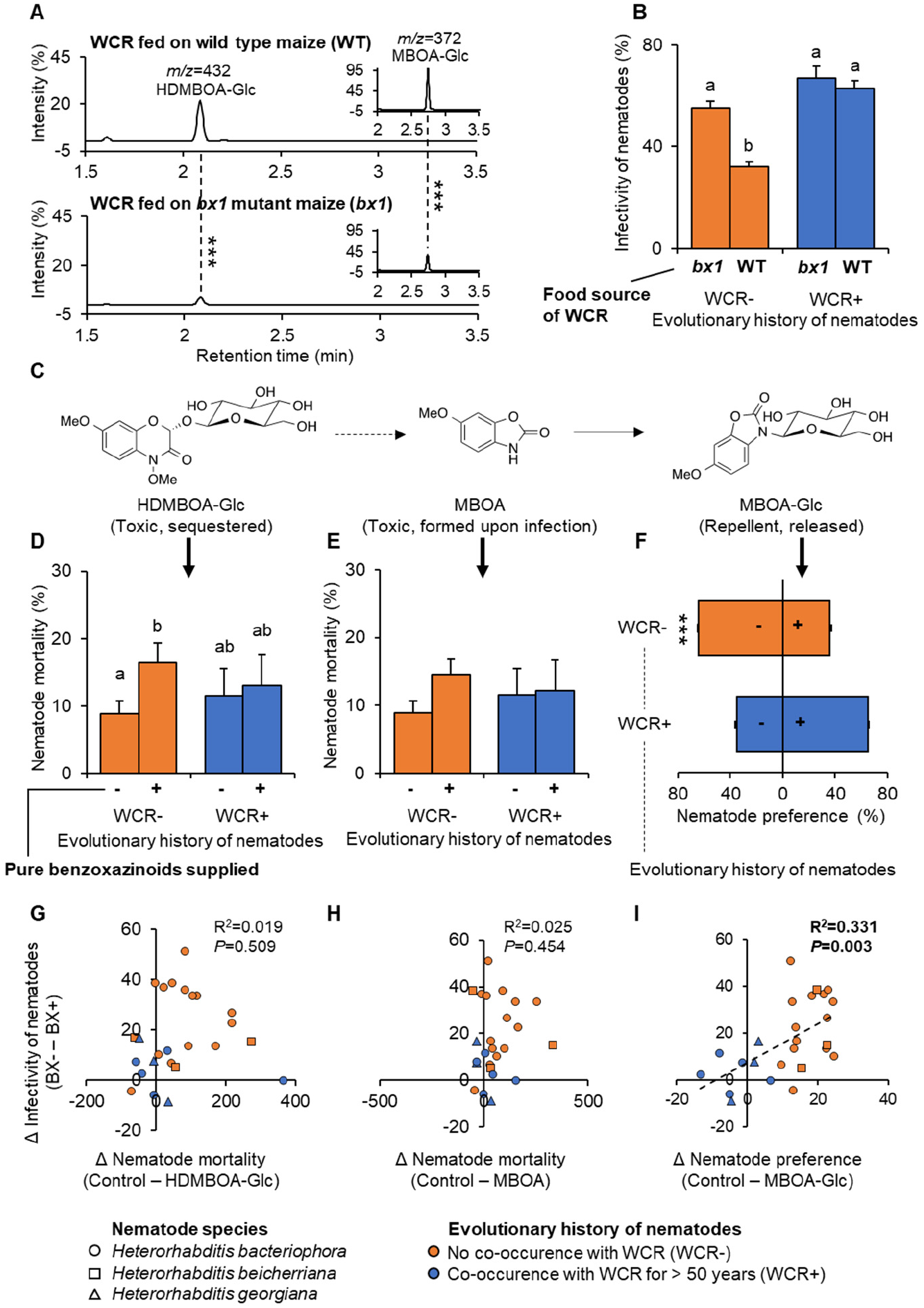
Nematodes from the primary range of the western corn rootworm are more resistant to sequestered benzoxazinoids. (**A**) Chromatograms of plant-derived benzoxazinoids in the body of western corn rootworm (WCR) larvae fed on wild type (WT, top) and benzoxazinoid-deficient *bx1* mutant maize plants (bottom). Asterisks indicate significant differences (*P*<0.001). (**B**) Infectivity of nematodes that share an evolutionary history with the western corn rootworm (WCR+, blue) or not (WCR-, orange) towards WCR larvae fed on WT or *bx1* mutant plants. Different letters indicate significant differences between treatments (False discovery rate corrected *P*<0.05). (**C**) Benzoxazinoids found in WCR larvae. Two-hydroxy-4,7-dimethoxy-1,4-benzoxazin-3-one *O*-glucoside (HDMBOA-Glc) accumulates in the larval body and is toxic for nematodes. Six-methoxy-2-benzoxazolinone (MBOA) is formed upon tissue disruption and nematode attack and is also toxic. Six-methoxy-2-benzoxazolinone *N*-glucoside (MBOA-Glc) is released by the larvae. It is not directly toxic but repels the nematodes. (**D-E**) Impact of physiologically relevant doses of HDMBOA-Glc (150 μg/mL) and MBOA (25 μg/mL) on nematode mortality. Different letters indicate significant differences between treatments (False discovery rate corrected *P*<0.05). (**F**) Impact of physiological doses of MBOA-Glc (3 μg/mL) on nematode attraction. Asterisks indicate a significant effect of MBOA-Glc (*P*<0.001). (**G-I**) Linear correlations between benzoxazinoid dependent infectivity (data from (**B**)) and in vitro benzoxazinoid resistance (data from (**D-F**)). R^2^ and *P*-values of linear regressions are shown. Dashed regression lines are shown for significant linear correlations.

Benzoxazinoids can protect the western corn rootworm from nematodes through a series of different, mutually non-exclusive mechanisms (Fig. 2C) (28). Six-methoxy-2-benzoxazolinone *N*-glucoside (MBOA-Glc) is an insect-specific conjugate formed from benzoxazinoid breakdown products that is released by the larvae and accumulates on their cuticule, thus making them less attractive to entomopathogenic nematodes. Two-hydroxy-4,7-dimethoxy-1,4-benzoxazin-3-one *O*-glucoside (HDMBOA-Glc) is contained within the larval body and is directly toxic to the nematodes (28). Upon nematode-infection, HDMBOA-Glc is broken down to 6-methoxy-2-benzoxazolinone (MBOA), which also reduces nematode survival (28). In choice experiments, MBOA-Glc reduced the attraction of nematodes that share no evolutionary history with the western corn rootworm, while nematodes from the primary range did not show a negative response towards MBOA-Glc (Fig. 2F and S6). Physiological doses of HDMBOA-Glc induced mortality in strains that shared no evolutionary history with the western corn rootworm, but not in strains from the primary range (Fig. 2D). No clear effects were found for MBOA-induced mortality (Fig 2E). Simple linear regression analysis revealed a positive correlation between the avoidance of MBOA-Glc and the benzoxazinoid-specific suppression of nematode infectivity (Fig. 2I). Model selection applied on a multiple linear regression including all explanatory factors and their interactions resulted in a model with the behavioral response towards MBOA-Glc alone explaining 61% of benzoxazinoid-dependent nematode infectivity across the different strains (Table S4). Taken together, these results show that nematodes from the primary range of the western corn rootworm do not avoid an insect-specific benzoxazinoid breakdown product and are more resistant to sequestered benzoxazinoids. Furthermore, the loss of behavioral avoidance of MBOA-Glc is associated with an increased capacity to infect benzoxazinoid-sequestering western corn rootworm larvae.

Benzoxazinoid tolerance in the nematode strains from maize fields of the primary range of the western corn rootworm may be confounded by population structure. Furthermore, their benzoxazinoid resistance may stem from exposure to benzoxazinoids exuded by maize roots rather than benzoxazinoids that are sequestered by the western corn rootworm. To test whether exposure to a benzoxazinoid-sequestering host can directly leads to the evolution of benzoxazinoid resistance, we designed a real-time evolution experiment. The benzoxazinoid-susceptible *H. bacteriophora* strain RW14 was multiplied and divided into 20 experimental (sub-)populations. Ten populations were then reared on banded cucumber beetle larvae and ten populations were reared on western corn rootworm larvae, both of which were fed on wild type maize roots. Using these two herbivore species allowed us to assess evolution on two natural hosts with different abilities to sequester benzoxazinoids. Subsequent infectivity tests were performed with wild-type and *bx1* fed western corn rootworm larvae to specifically assess the evolution of benzoxazinoid-resistance following exposure to the different herbivores. In the F1 generation, all experimental nematode populations showed reduced infectiveness towards wild type-fed western corn rootworm larvae compared to *bx1*-fed larvae (Fig 3A and S7). After five generations of selection, nematodes reared on banded cucumber beetle larvae still showed reduced infectivity towards wild-type fed western corn rootworm larvae, even though the effect was less strong than in the F1 generation (Fig. 3B and S8). By contrast, nematodes reared on the western corn rootworm were overall better able to infect the western corn rootworm and did no longer show a reduction in infectivity on wild type fed larvae compared to *bx1*-fed larvae (Fig. 3B and S8). The nematodes were even more successful on wild type fed western corn rootworm larvae than on *bx1*-mutant fed larvae (Fig. 3B). Thus, selection on a benzoxazinoid-sequestering host over five generations is sufficient for susceptible nematodes to evolve complete resistance to benzoxazinoid-dependent defenses.

**Fig. 3.**
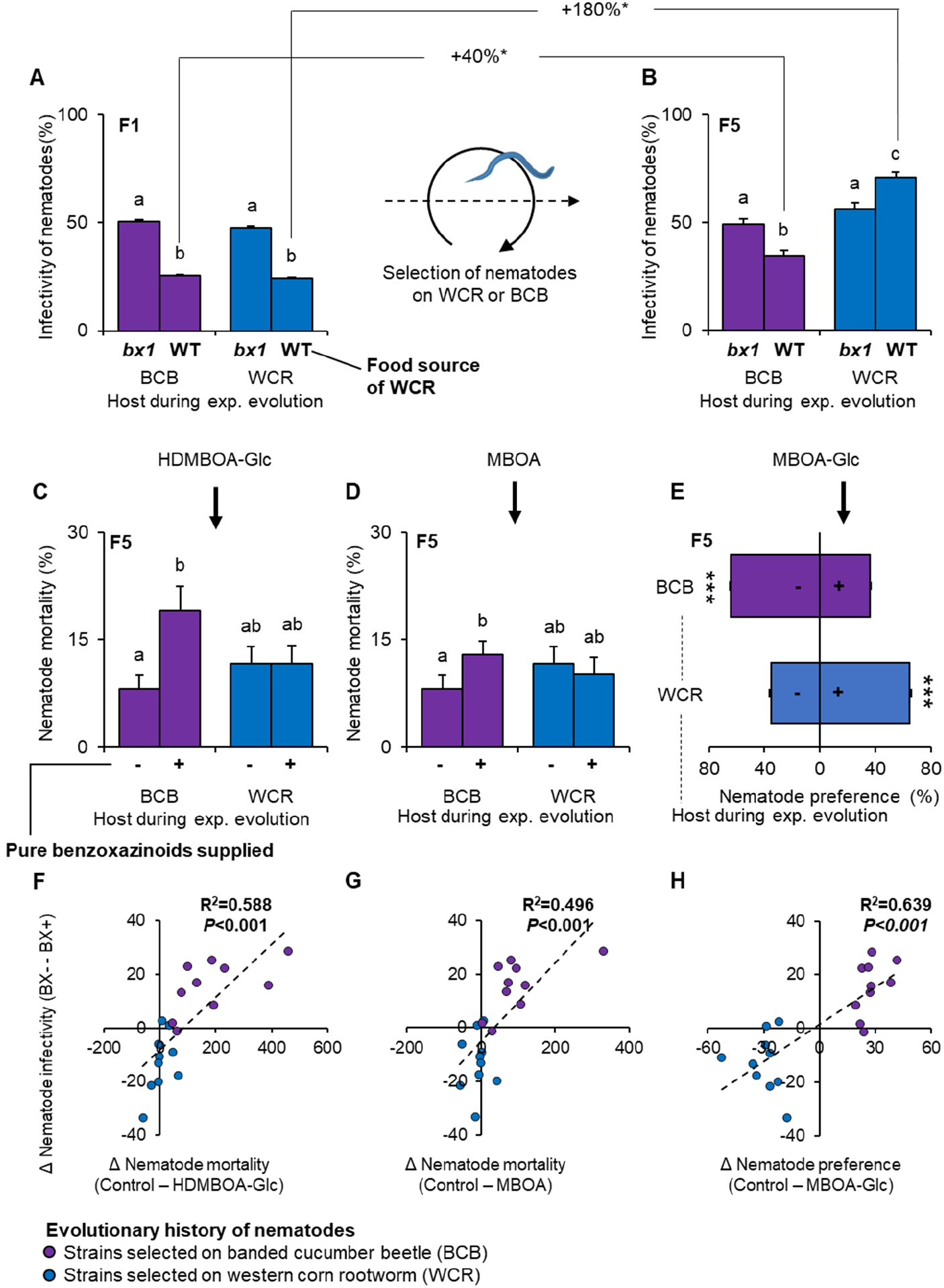
Rapid evolution of benzoxazinoid resistance and infectivity in nematodes exposed to the western corn rootworm. (**A**) Infectivity of F1 nematodes reared on the western corn rootworm (WCR+, blue) or the banded cucumber beetle (BCB, purple) for one generation, exposed to WCR larvae fed on WT or *bx1* mutant plants. Different letters indicate significant differences between treatments (False discovery rate corrected least square means, *P*<0.05). (**B**) Infectivity of F5 nematodes reared on the western corn rootworm (WCR+, blue) or the banded cucumber beetle (BCB, purple) for five generations and exposed to WCR larvae fed on WT or *bx1* mutant plants. Different letters indicate significant differences between treatments (False discovery rate corrected *P*<0.05). Significant changes (*P*<0.05) in infectivity between F1 and F5 nematodes are indicated by asterisks. (**C-D**) Impact of physiologically relevant doses of HDMBOA-Glc (150 μg/mL) and MBOA (25 μg/mL) on nematode mortality. Different letters indicate significant differences between treatments (False discovery rate corrected *P*<0.05). (**E**) Impact of physiological doses of MBOA-Glc (3 μg/mL) on nematode attraction. Asterisks indicate a significant effect of MBOA-Glc (*P*<0.001). (**F-H**) Linear correlations between benzoxazinoid dependent infectivity (data from (**B**)) and in vitro benzoxazinoid resistance (data from (**C-E**)). R^2^ and *P*-values of linear regressions are shown. Dashed regression lines are drawn for significant correlations.

To understand how the western corn rootworm-selected nematode populations may achieve higher infectiveness towards benzoxazinoid-containing western corn rootworm larvae, we subjected them to the same series of bioassays as the natural strain collection before (Fig. 3 C-E). Nematodes selected on banded cucumber beetle larvae preferred to avoid MBOA-Glc, while nematodes selected on western corn rootworm larvae were attracted by this compound (Fig. 3E and S9). Furthermore, HDMBOA-Glc and MBOA reduced the survival of nematodes selected on the banded cucumber beetle, but did not reduce survival of nematodes selected on the western corn rootworm (Fig. 3C-D and S9). Simple linear regression analysis revealed a correlation between the behavioral response to MBOA-Glc and benzoxazinoid-specific nematode infectivity as well as HDMBOA-Glc and MBOA toxicity and benzoxazinoid-specific nematode infectivity (Fig. 3 F-H). Multiple linear regression including all explanatory factors and their interactions, followed by model selection and averaging, resulted in a model with MBOA-Glc repellence and HDMBOA-Glc toxicity explaining 82% of the benzoxazinoid-dependent nematode infectivity across strains (Table S4). Thus, similar to the pattern observed for the natural strains, nematodes exposed to the western corn rootworm for five generations do no longer avoid an insect-specific benzoxazinoid breakdown product and become resistant to a sequestered benzoxazinoid and its breakdown product. Furthermore, their increased capacity to infect benzoxazinoid-sequestering western corn rootworm larvae is associated with both a loss of behavioral aversion and reduced benzoxazinoid toxicity.

## Discussion

Herbivore natural enemies are often exposed to plant defense metabolites, either by coming into contact with plants directly or by preying on herbivores that contain plant defenses (33–36). Many herbivore natural enemies have been found to avoid plant defenses by rejecting herbivores that accumulate or sequester toxins (37–39). However, avoidance comes with significant costs, especially when mobility and host availability are limited. Soil-borne herbivore natural enemies such as entomopathogenic nematodes can typically only cover short distances (40) and may have a limited choice of hosts in agricultural environments. Thus, they are likely to be under considerable pressure to overcome, rather than to avoid, plant toxins, which may explain why nematodes that share an evolutionary history with the western corn rootworm show benzoxazinoid resistance in the form of a loss of behavioral aversion. Together with their increased capacity to withstand the toxic effects of benzoxazinoids, this behavioral shift likely allows the nematodes to prey successfully on benzoxazinoid-sequestering hosts such as the western corn rootworm. The exact contribution of the western corn rootworm to nematode evolution in the field relative to other soil-borne organisms that are commonly present in US maize fields and may modulate benzoxazinoid exposure requires further study. Given the dominance of the western corn rootworm in maize fields in the areas from which the benzoxazinoid-resistant nematodes were sampled, however, we consider it likely that their resistance traits originate from the selection pressure exerted by the western corn rootworm.

The real-time evolution experiment further demonstrates that benzoxazinoid resistance in nematodes can evolve rapidly following exposure to the western corn rootworm, most likely through standing genetic variation. The finding that nematodes selected on the banded cucumber beetle also become slightly less susceptible to benzoxazinoid-dependent defenses is noteworthy in this context, as it may indicate that plant defense resistance in herbivore natural enemies can also evolve in the absence of a defense-sequestering herbivore, for instance through exposure to residual plant defense levels in the soil as well as the gut and the frass of the herbivore. Whether evolved resistance to plant toxins is common in herbivore natural enemies remains to be determined. Given that exposure of natural enemies to plant toxins is frequent in nature (33–36) and that different natural enemies have been reported to resist and accumulate plant toxins (18–21), we expect evolved behavioral avoidance, metabolic resistance and tolerance strategies to be widespread among members of the third trophic level. Host diversity and diet breadth will likely determine the prevalence and biochemical architecture of these traits in herbivore natural enemies (20).

The diversity of plant defense metabolites and arthropod herbivores in nature is thought to be the result of an ongoing co-evolutionary arms race (41–45). Herbivore natural enemies have been shown to be negatively affected by plant defense metabolites in this context (46–48). The possibility that herbivore natural enemies may evolve resistance to increasingly toxic herbivores been considered (17,18), but, to the best of our knowledge, not been addressed through manipulative experiments. Instead, adaptations of herbivore natural enemies to plant chemicals have been investigated in detail in the context of plant volatiles that serve as foraging cues for herbivore natural enemies (48–50), including volatiles that attracted entomopathogenic nematodes (51), and in the context of extrafloral nectar as food source (50, 52). Although conducted in an agricultural system, the present study does support an evolutionary link between the co-evolutionary arms race of plants and herbivores and the third trophic level by showing that plant defense metabolites may influence the evolution of herbivore natural enemies as they are transferred through adapted herbivores. Trophic transfer of defense metabolites may promote the specialization and diversification of herbivore natural enemies (46). In wild systems, resistance to plant defenses by herbivore natural enemies may reduce the penalty for plants facing adapted herbivores (17), which again may reduce the negative selection pressure on basal plant defense metabolites and thereby contribute to within-plant chemical diversity. Detailed mechanistic and evolutionary studies, including broad phylogenetic analyses of multitrophic interaction networks in natural systems, could help to shed further light on these hypotheses.

Reducing the use of synthetic pesticides is an important aim in sustainable agriculture. The use of herbivore natural enemies as biological control agents is a promising strategy in this context (53). However, the efficacy of biocontrol agents is often limited, and a better understanding of the factors that determine their success is thus important to improve their use (54). Our study confirms that plant toxins can limit the capacity of natural enemies to control agricultural pests (17, 28), but also shows that plant toxin susceptibility shows pronounced heritable variation and can be offset rapidly through artificial selection. Plant toxin resistance thus represents a promising breeding target for the improvement of biological control agents.

## Materials and Methods

### Insects

Western corn rootworm (WCR, *Diabrotica virgifera virgifera* LeConte) eggs were supplied by the USDA-ARS-NCARL, Brookings, SD, USA. Banded cucumber beetle (BCB, *D. balteata* LeConte) eggs were obtained from Syngenta Crop Protection AG, Stein, CH. After hatching, WCR and BCB neonates were reared on freshly germinated wild type (B73) maize seeds or on the benzoxazinoid mutant line *bx1* (55). Third instar WCR and BCB larvae were used for all experiments. *Galleria mellonella* larvae were bought from Fischereibedarf Wenger, Bern, CH, and maintained at 8°C until use for nematode rearing.

### Nematodes

#### Nematode collection, identification and rearing

We established a collection of 25 different *Heterorhabditis* spp. Detailed information on the different strains can be found in Table S1. Strains were either obtained from collaborators or collected from the field. Field collections were realized by collecting 45 soil cores (10 cm depth, 2 cm diameter) for each individual location. Soil cores were pooled for each location, homogenized and separated into 20 plastic containers (250 mL, Semadeni AG, CH). Five *G. mellonella* larvae were then placed on the soil surface of each container. The containers were closed with a plastic lid and incubated in darkness under 24 ± 2 °C. Five to ten days later, all nematode-infected *G. mellonella* larvae were individually transferred to white traps (56), and emerging nematodes were used to infect another set of *G. mellonella* larvae (57). Irrespective of whether they were isolated from the field or obtained from collaborators, all nematodes were identified by internal transcribed spacer rRNA gene sequencing as described previously (58–60). The collected nematode progeny was maintained in 250 ml flasks (Greiner Bio-One GmbH, Frickenhausen, DE) at a density of one EPN per microliter tap water at 10 °C. Nematode strains were refreshed by multiplying them on *G. mellonella* larvae every 2-3 months. Nematodes that were less than one-month old were used for all experiments.

### Benzoxazinoid analyses

Benzoxazinoids from BCB larvae and WCR larvae fed on B73 or *bx1* maize plants were extracted in 50% MeOH + 50% H2O+ 0.5% formic acid. Five flash frozen larvae (~40 mg) were pooled and ground in 400 μL extraction buffer, and 5 replicates (each consisting of a pool of 5 larvae) were analyzed for each species and food source. The extracts were vortexed for 1 min and centrifuged twice at 17,000 g, at 4 °C. The supernatants were then analyzed on an Acquity UHPLC-MS system equipped with an electrospray source (Waters i-Class UHPLC-QDA, USA). The method was modified from the one described previously (10). Briefly, the elution profile was: 0–3.5 min, 99– 72.5% A in B; 3.5–5.5 min, 100% B; 5.5–7.5 min 99% A in B. The injection volume was 1 μL. DIMBOA, MBOA, DIMBOA-Glc, HDMBOA-Glc and MBOA-Glc were all quantified in ESI-using selective ion recording (SIR) and external standard curves.

### Real-time evolution

An experimental selection experiment was carried out using the RW14 nematode strain. A batch of newly hatched RW14 nematodes was aliquoted into 20 sub-populations, each consisting of 20000 nematodes. Half the sub-populations were then reared on WCR larvae, and the other half on BCB larvae. Infection was performed in solo cups as described below in the section “Nematode infectivity”. For each population, 30 solo cups containing 5 larvae each were infected with 500 nematodes. Five days later, all the infected larvae of the same sub-population were collected together and transferred to white traps for collecting nematode progenies. In between each generation of selection, the populations were amplified in *G. mellonella* larvae by infecting 15 larvae per population with 200 nematodes each. The different populations were selected on WCR and BCB larvae for a total of 10 generations (5 generations within the selection host and 5 amplification steps in between), and F1 and F5 nematodes (referring to the number of generations on the selection host) of 10 independently selected sub-populations were phenotyped.

### Nematode infectivity

To quantify the infectivity of the nematodes, three to five WCR or BCB larvae were placed into individual solo cups (30 mL, Frontier Scientific Services, Inc., DE) containing a 5 mm layer of moist, autoclaved sand (Selmaterra, Bigler Samen AG, Thun, CH). Five hundred nematodes in 500 μL tap water were applied into each solo-cup. After incubating the cups at 28 ± 0.5 °C for 5 days, the percentage of nematode-infected larvae in each solo-cup was determined. Larvae were reared on either wild type (B73) or *bx1* mutant plants. Each solo cup was treated as an independent replicate. The exact numbers of independent biological replicates for the individual experiments and treatments are provided in Table S2.

### Nematode behavior

To test the effect of MBOA-Glc on nematode behavior, we used the approach developed earlier (28). Briefly, a 5 mm agarose (5 g/L, Sigma Aldrich Chemie, CHE) layer was poured into a Petridish (9 cm diameter, Greiner Bio-One GmbH, Frickenhausen, DE). Three 5-mm-diameter wells were created equidistantly in the agar layer: one in the center of the plate, one on the right and one on the left side of the plate. One hundred nematodes in 100 μL of water were dispensed the central well. The remaining two wells were filled with either 50 μL BCB exudates and 50 μL tap water or with 50 μL BCB exudates + 50 μL MBOA-Glc (3 ng/μL in tap water). BCB exudates were used to elicit nematode search behavior. Exudates were obtained by rinsing third instar BCB larva with 50 μL tap water. Nematode preference was recorded 24 h after the start of the experiment by counting the number of nematodes within different sectors of the Petri dishes. The dishes were divided into 4 sectors of equal size, with two opposite sectors containing the treatment wells. Nematodes within the treatment sectors were counted. Each petri dish was treated as independent biological replicates, and 18-20 replicates were carried out for the different experiments and treatments (Table S2).

### Nematode performance

The effects of benzoxazinoids on nematode survival and infectivity were tested as described (28). Briefly, 4000 nematodes were incubated in 4 mL tap water containing either MBOA (25 μg/mL) or HDMBOA-Glc (150 μg/mL). These concentrations represent physiologically relevant doses (28). Nematodes were kept in 50 mL flasks (Greiner Bio-One GmbH, Frickenhausen, DE) and incubated at 28°C. Nematodes incubated in tap water were used as controls. The number of dead and living nematodes was recorded 7 days after incubation. Flasks were treated as independent biological replicates, and 8-10 replicates were carried out for the different experiments and treatments (Table S2).

### Statistical analyses

Generalized linear mixed models (GLMMs; distribution: binomial, link function: logit) were used to analyze mortality, infectivity and preference bioassays. Effects of nematode origin / experimental evolution host, different treatments and their interactions were used as main factors, and nematode strains / populations were used as a random factor. Wald tests were performed to assess significance of treatment effects. Pairwise comparisons of Estimated Marginal Means (EMMeans) corrected by the False Discovery Rate (FDR) method were used as post hoc tests (61). To determine correlations between benzoxazinoid resistance traits and benzoxazinoid dependent infectivity of the different nematode strains and populations the delta between BX+ (wild type fed WCR larvae) and BX- (*bx1*-mutant fed larvae) infectivity (i.e. benzoxazinoid dependent infectivity) was calculated based on EMMeans. Then, different types of benzoxazinoid resistance were determined for each strain by calculating the respective deltas as well. Linear models were then employed using benzoxazinoid dependent infectivity as response variable and the different resistance deltas as explanatory variables. A complementary approach to test for multiple explanatory variables and their interactions was executed using model selection on a multiple linear regression including all resistance deltas and their interactions as explanatory variables. The Aikake Information Criterion corrected for small samples sizes (AICc) was used to rank submodels. Models showing a delta-AICc ≤ 2 were combined using a model averaging procedure, which allows computing the relative importance of each resistance delta in the averaged model. The packages used for the different analyses were “RVAideMemoire”, “car”, “emmeans”, “MASS”, “lme4”, “lmerTest” and “MuMIn” (62). All the analyses were conducted in R 3.5.1.

## Acknowledgements

We thank Ralf-Udo Ehlers (e-nema GmbH, Germany) for sharing nematode strains, Fausto Prada, Lisa Thönen, Virginia Hill, Anja Boss and Wei Huang for their assistance with laboratory experiments, David Ermacora for nematode rearing and Anouk Guyer, Zixiao Zhao and various interns from the Hibbard laboratory for field assistance. We also thank Lance Meinke (University of Nebraska), Joe Spencer (Illinois Natural History Survey), Sarah Zukoff (Kansas State University), Ken Ostlie (University of Minnesota), and Billy Fuller & Brad McMannus (South Dakota State University) for identifying fields for nematode collection.

## Author contributions

M.E. and C.A.M.R. conceived the original project. M.E., C.A.M.R. and R.A.R.M. designed experiments. M.E., C.A.M.R. R.A.R.M., B.H. and C.P. supervised experiments. X.Z., C.C.M.A, R.A.R.M, D.v.C., S.G. and L.H. performed experiments. M.E., R.A.R.M, X.Z. C.P and S.G. analyzed experiments. M.E. wrote the first draft of the manuscript. All authors contributed to the final version of the manuscript.

## Data and materials availability

Some of the nematode strains were obtained under a materials transfer agreement. The data generated for this manuscript will be made available through a public repository at a later stage.

---

**Fig. S1.**
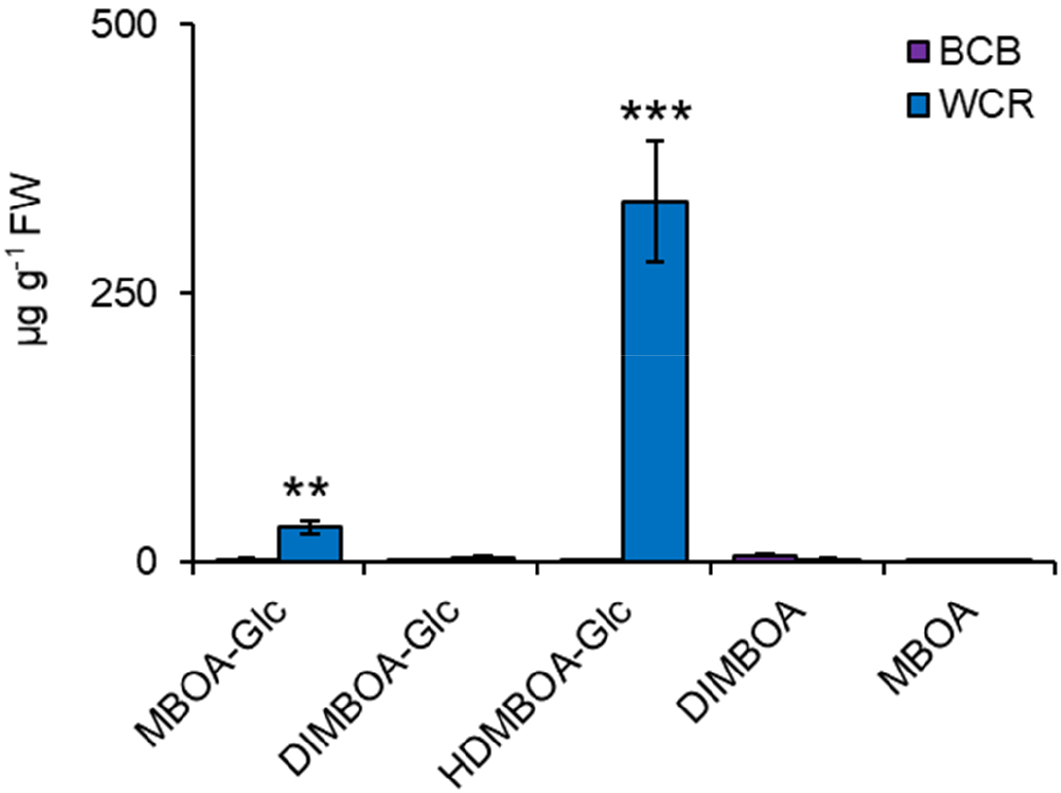
Quantification of plant-derived benzoxazinoids in different nematode hosts. Absolute quantities of benzoxazinoids extracted from larvae of the western corn rootworm (WCR; blue) and the banded cucumber beetle (BCB) are shown. Asterisks indicate significant differences between herbivore species (\*\**P*<0.01; *** *P* <0.001).

**Fig. S2.**
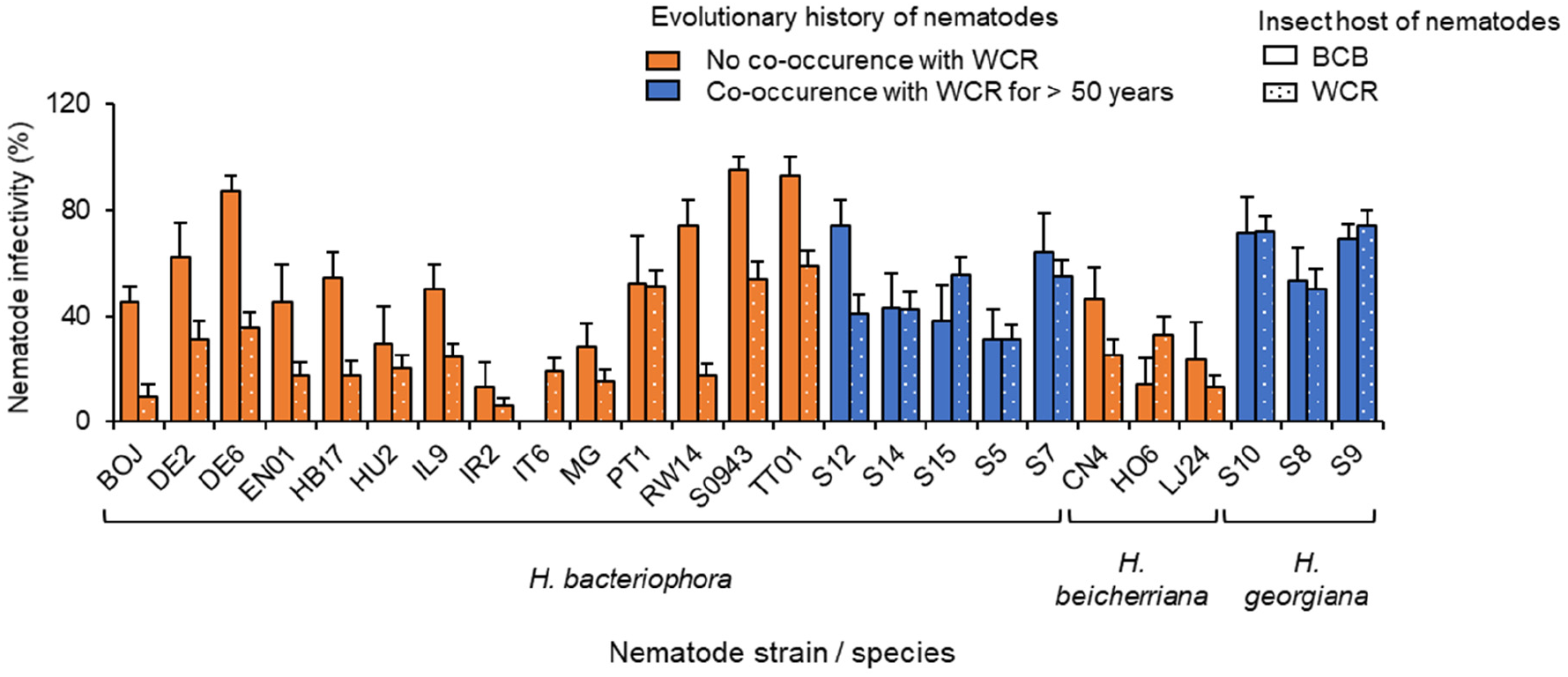
Infectivity of individual nematode strains towards different herbivores. Infectivity of individual nematode strains from the primary range of the western corn rootworm (blue) and other parts of the world (orange) towards the western corn rootworm (WCR; plain filled bars) and the banded cucumber beetle (BCB; dotted bars) is shown.

**Fig. S3.**
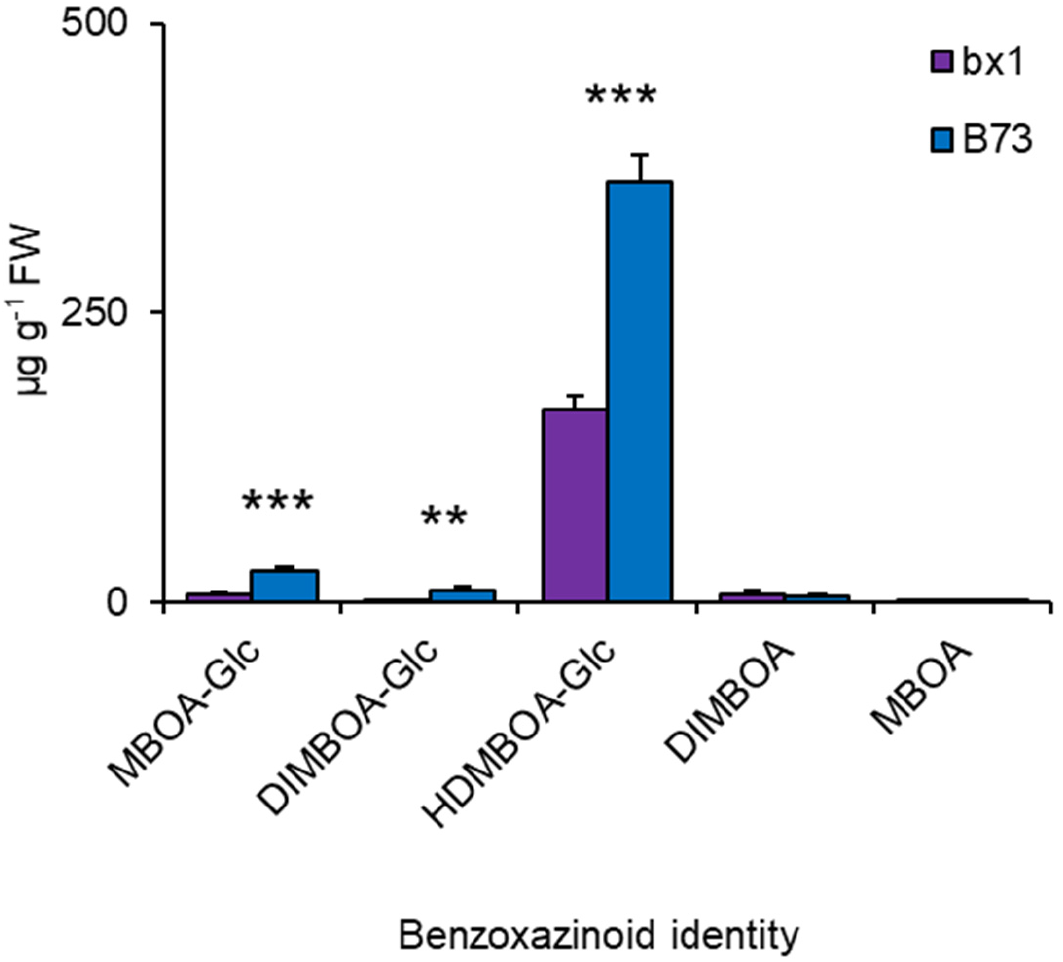
Quantification of benzoxazinoids in western corn rootworm larvae fed on wild type and *bx1* mutant maize roots. Absolute quantities of benzoxazinoids extracted from larvae of the western corn rootworm fed on *bx1* mutant (*bx1*) and wild type (WT) maize roots are shown. Asterisks indicate significant differences between plant genotypes (\*\**P* <0.01; *** *P* <0.001).

**Fig. S4.**
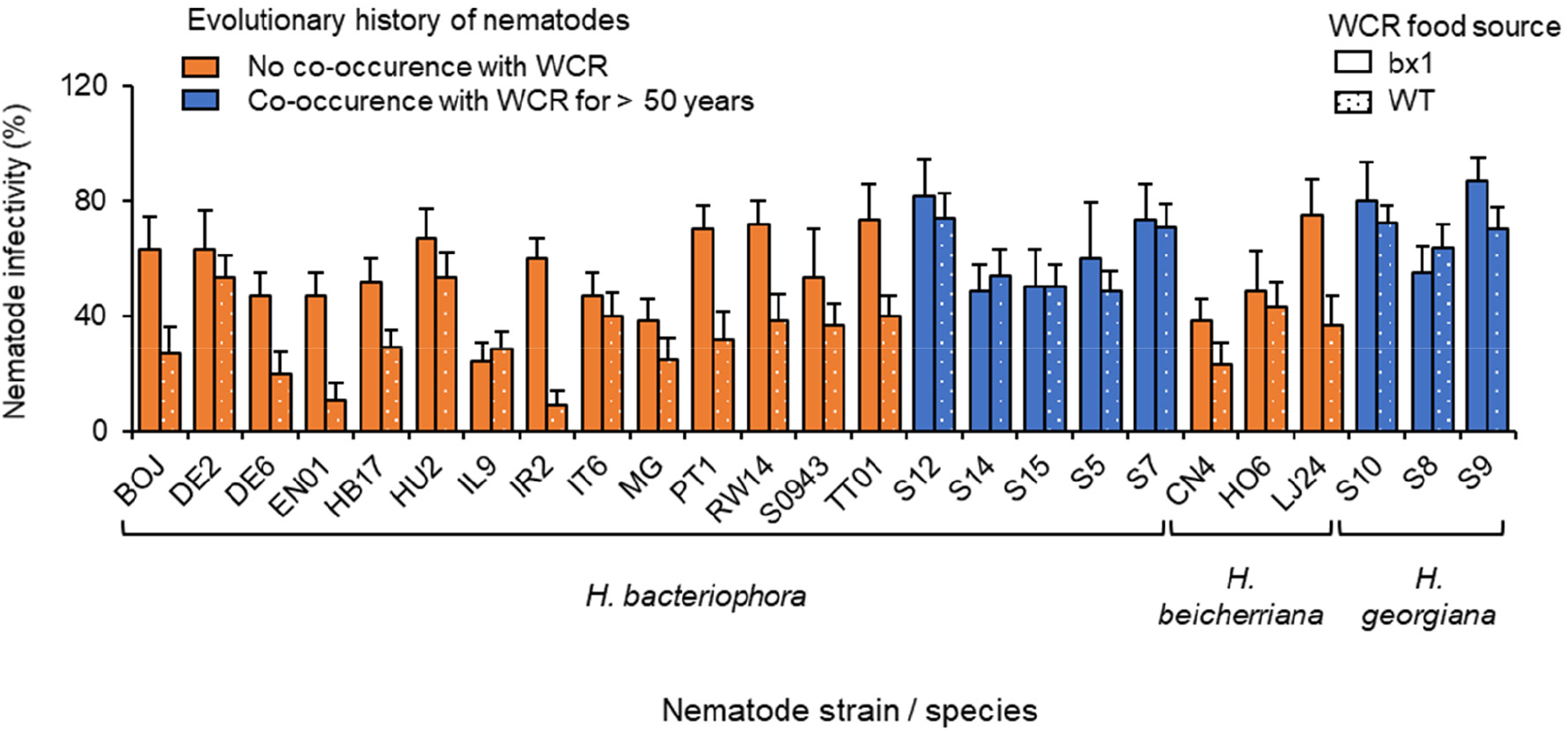
Impact of herbivore-sequestered benzoxazinoids on the infectivity of individual nematode strains. Infectivity of individual nematode strains from the primary range of the western corn rootworm (WCR; blue) and other parts of the world (orange) towards WCR larvae fed on *bx1* mutant (*bx1*; plain filled bars) and wild type (WT; dotted bars) maize roots is shown.

**Fig. S5.**
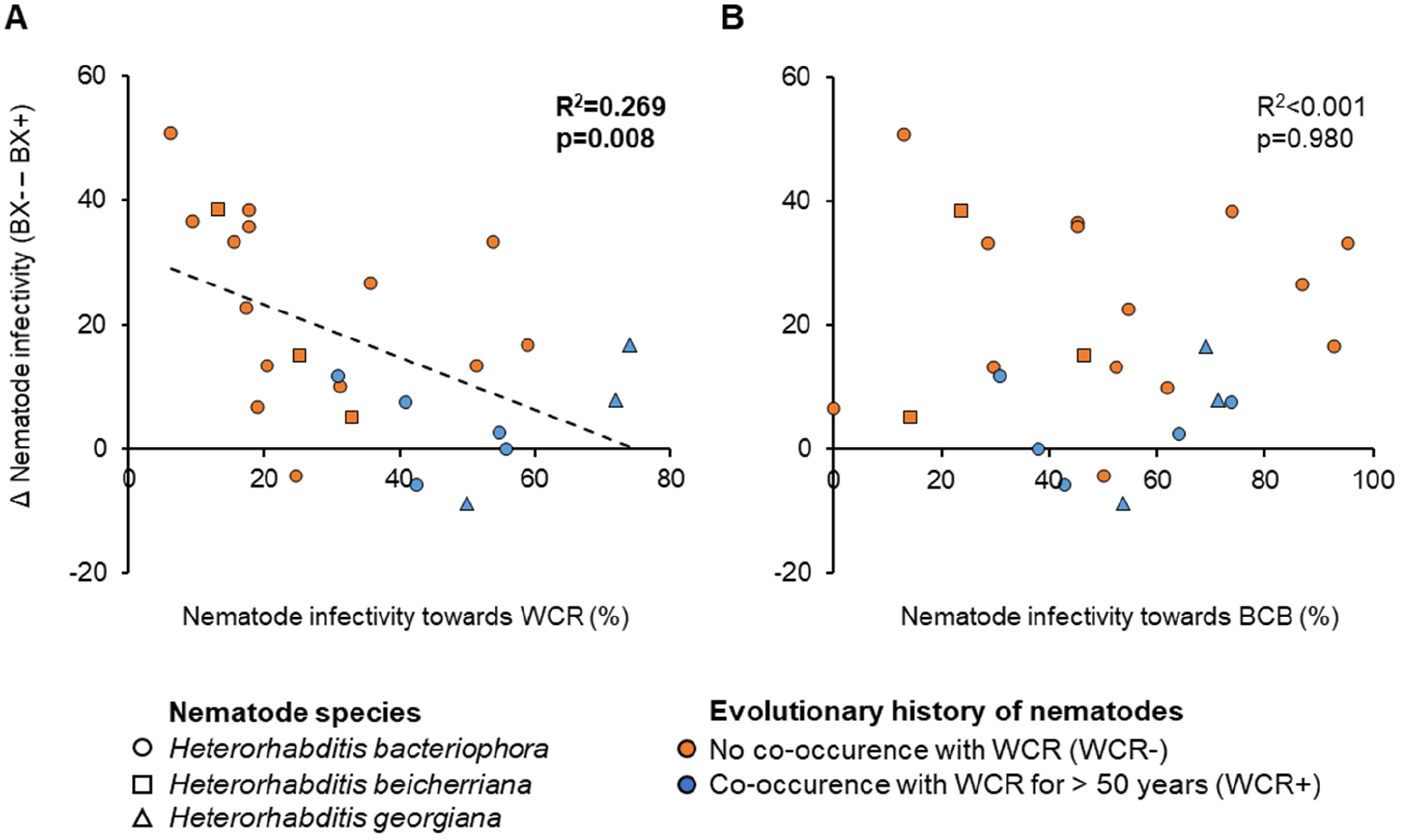
Correlations between benzoxazinoid-dependent infectivity and infectivity towards different herbivores. Correlations are shown between the capacity of the different nematode strains to infect western corn rootworm (WCR) and banded cucumber beetle (BCB) larvae (Fig. S2) and their capacity to withstand sequestered benzoxazinoids (Fig. S4). Benzoxazinoid resistance is calculated by taking the difference in infectivity of the individual strains between *bx1* mutant (BX-) and wild type (BX+) maize root fed western corn rootworm larvae. Positive values correspond to higher infectiveness towards *bx1* mutant fed larvae. Significant correlations are indicated with dashed lines. R^2^ and *P*-values from Pearson product-moment correlations are provided.

**Fig. S6.**
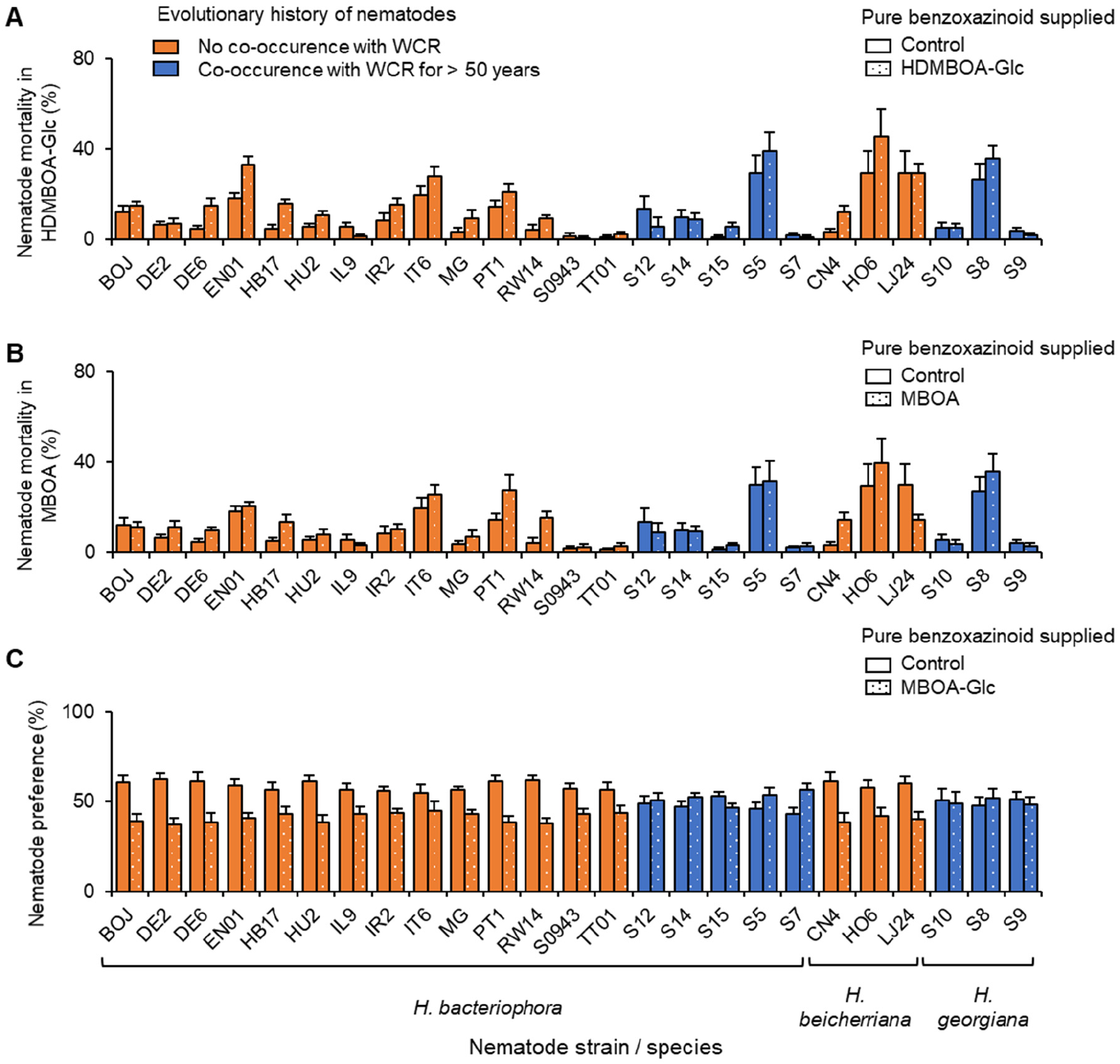
Impact of pure benzoxazinoids on individual nematode strains. **(A)** Mortality of individual nematode strains from the primary range of the western corn rootworm (WCR; blue) and other parts of the world (orange) treated with water (plain filled bars) or 150 μg/mL HDMBOA-Glc (dotted bars) is shown. **(B)** Mortality of nematodes treated with water or 25 μg/mL MBOA. **(C)** Preference of nematodes for water or 3 μg/mL MBOA-Glc.

**Fig. S7.**
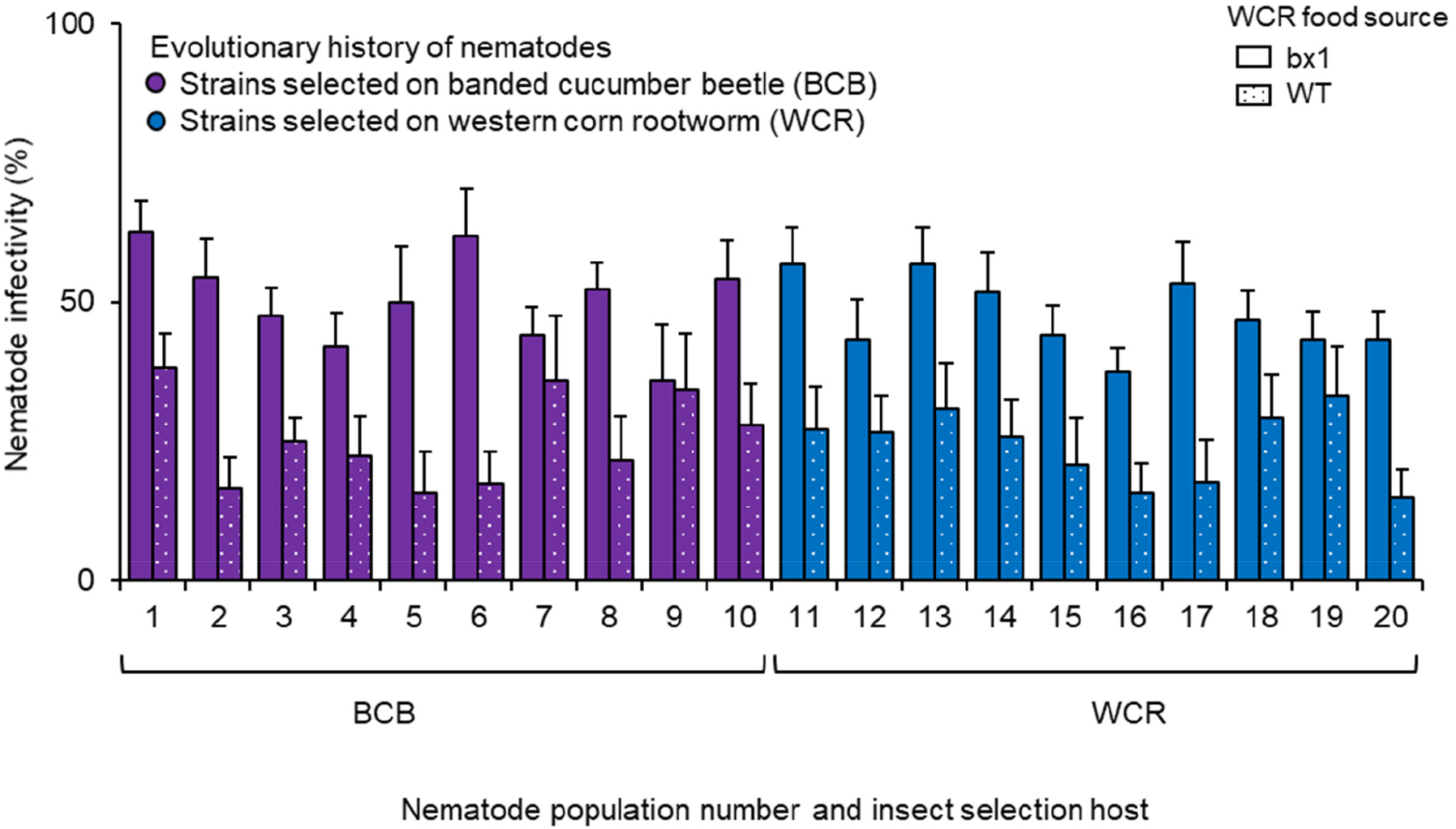
Impact of herbivore-sequestered benzoxazinoids on nematode infectivity after one generation of artificial selection on different herbivores. Infectivity of individual nematode strains selected on the western corn rootworm (WCR; blue) or the banded cucumber beetle (BCB; purple) towards WCR larvae fed on *bx1* mutant (*bx1*; plain filled bars) and wild type (WT; dotted bars) maize roots is shown.

**Fig. S8.**
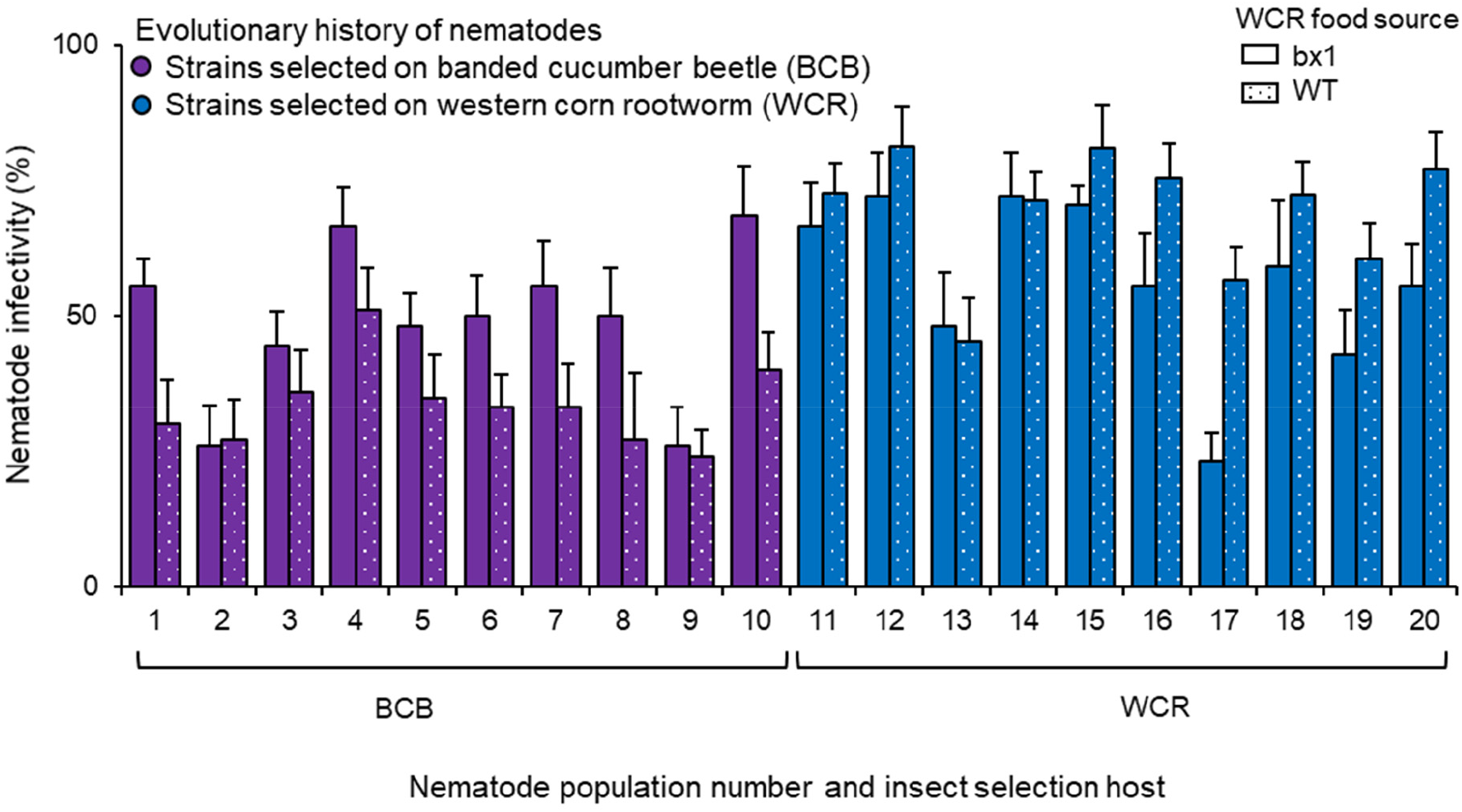
Impact of herbivore-sequestered benzoxazinoids on nematode infectivity after five generations of artificial selection on different herbivores. Infectivity of individual nematode strains selected on the western corn rootworm (WCR; blue) or the banded cucumber beetle (BCB; purple) towards WCR larvae fed on *bx1* mutant (*bx1*; plain filled bars) and wild type (WT; dotted bars) maize roots is shown.

**Fig. S9.**
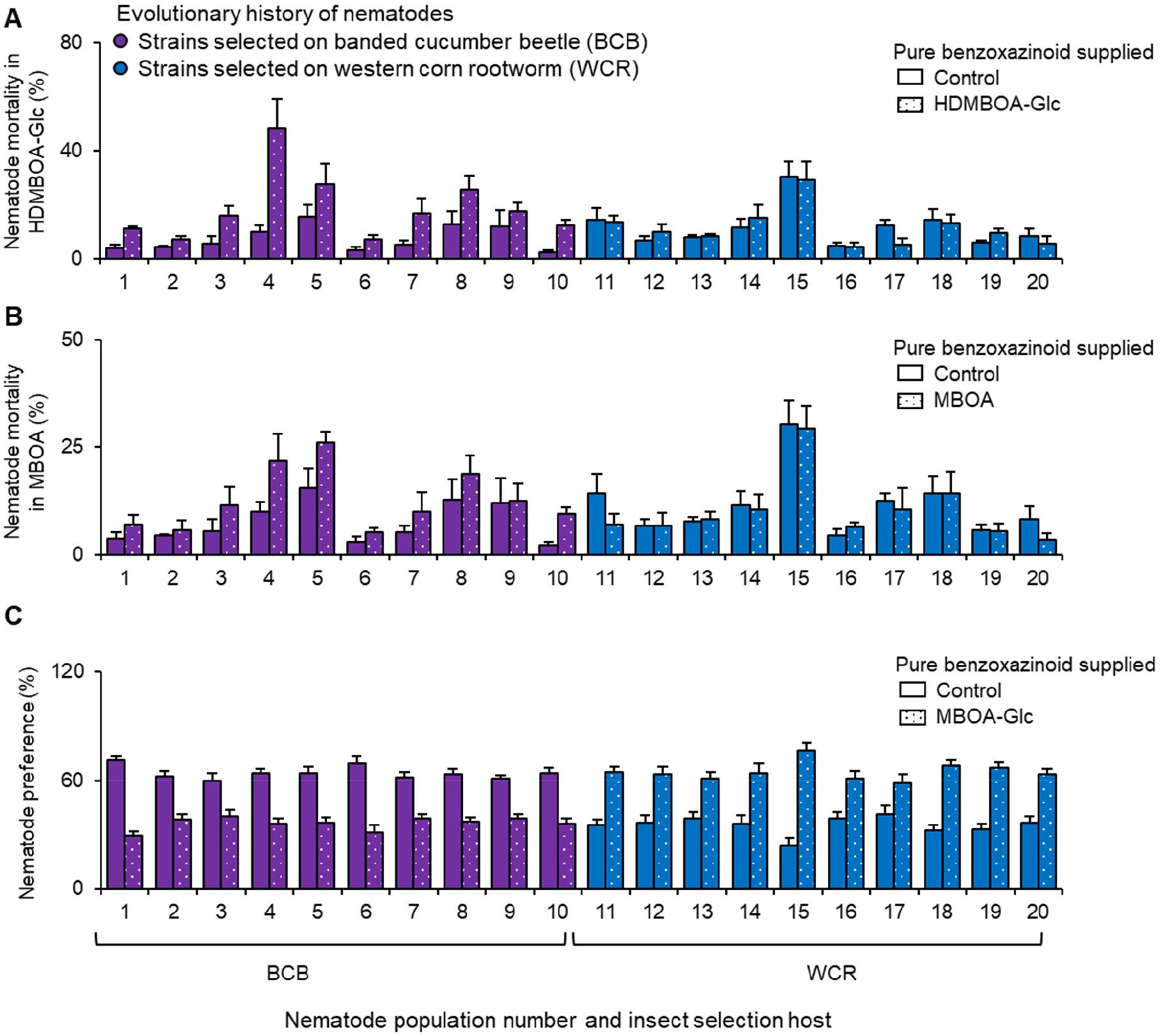
Impact of pure benzoxazinoids on individual nematode populations after 5 generations of artificial selection. **(A)** Mortality of individual nematode strains after five generations of selection on the western corn rootworm (WCR; blue) or the banded cucumber beetle (BCB; purple) treated with water (plain filled bars) or 150 μg/mL HDMBOA-Glc (dotted bars) is shown. **(B)** Mortality of nematodes treated with water or 25 μg/mL MBOA. **(C)** Preference of nematodes for water or 3 μg/mL MBOA-Glc.

**Table S1.**
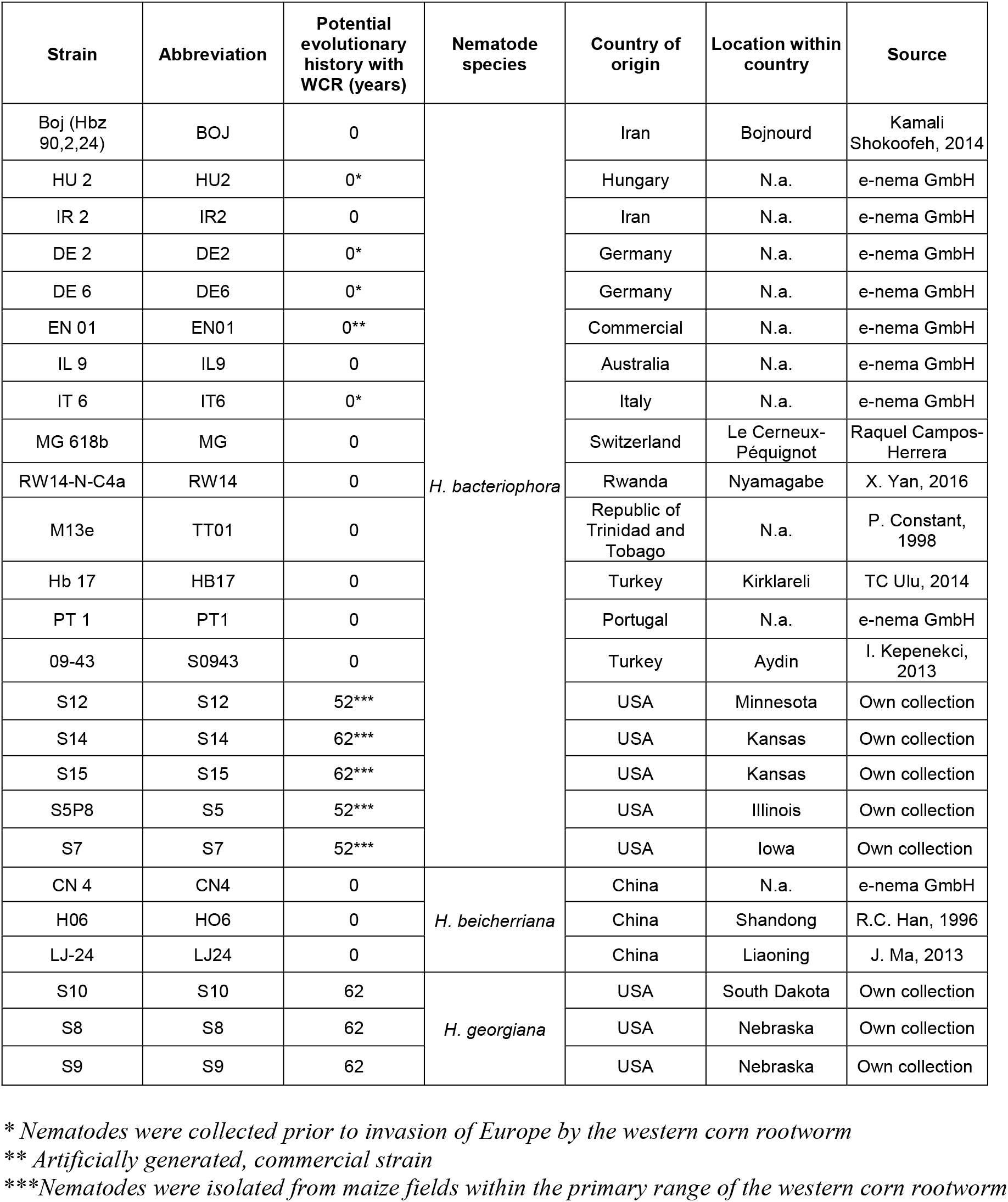
Source and evolutionary histories of the different nematode strains

| Strain | Abbreviation | Potential evolutionary history with WCR (years) | Nematode species | Country of origin | Location within country | Source |
| --- | --- | --- | --- | --- | --- | --- |
| Boj (Hbz 90,2,24) | BOJ | 0 | *H. bacteriophora* | Iran | Bojnourd | Kamali Shokoofeh, 2014 |
| HU 2 | HU2 | 0* |  | Hungary | N.a. | e-nema GmbH |
| IR 2 | IR2 | 0 |  | Iran | N.a. | e-nema GmbH |
| DE 2 | DE2 | 0* |  | Germany | N.a. | e-nema GmbH |
| DE 6 | DE6 | 0* |  | Germany | N.a. | e-nema GmbH |
| EN 01 | EN01 | 0** |  | Commercial | N.a. | e-nema GmbH |
| IL 9 | IL9 | 0 |  | Australia | N.a. | e-nema GmbH |
| IT 6 | IT6 | 0* |  | Italy | N.a. | e-nema GmbH |
| MG 618b | MG | 0 |  | Switzerland | Le Cerneux-Péquignot | Raquel Campos-Herrera |
| RW14-N-C4a | RW14 | 0 |  | Rwanda | Nyamagabe | X. Yan, 2016 |
| M13e | TT01 | 0 |  | Republic of Trinidad and Tobago | N.a. | P. Constant, 1998 |
| Hb 17 | HB17 | 0 |  | Turkey | Kirklareli | TC Ulu, 2014 |
| PT 1 | PT1 | 0 |  | Portugal | N.a. | e-nema GmbH |
| 09-43 | S0943 | 0 |  | Turkey | Aydin | I. Kepenekci, 2013 |
| S12 | S12 | 52*** |  | USA | Minnesota | Own collection |
| S14 | S14 | 62*** |  | USA | Kansas | Own collection |
| S15 | S15 | 62*** |  | USA | Kansas | Own collection |
| S5P8 | S5 | 52*** |  | USA | Illinois | Own collection |
| S7 | S7 | 52*** |  | USA | Iowa | Own collection |
| CN 4 | CN4 | 0 | *H. beicherriana* | China | N.a. | e-nema GmbH |
| H06 | HO6 | 0 |  | China | Shandong | R.C. Han, 1996 |
| LJ-24 | LJ24 | 0 |  | China | Liaoning | J. Ma, 2013 |
| S10 | S10 | 62 | *H. georgiana* | USA | South Dakota | Own collection |
| S8 | S8 | 62 |  | USA | Nebraska | Own collection |
| S9 | S9 | 62 |  | USA | Nebraska | Own collection |

\* Nematodes were collected prior to invasion of Europe by the western corn rootworm
\*\* Artificially generated, commercial strain
\*\*\* Nematodes were isolated from maize fields within the primary range of the western corn rootworm

**Table S2.**
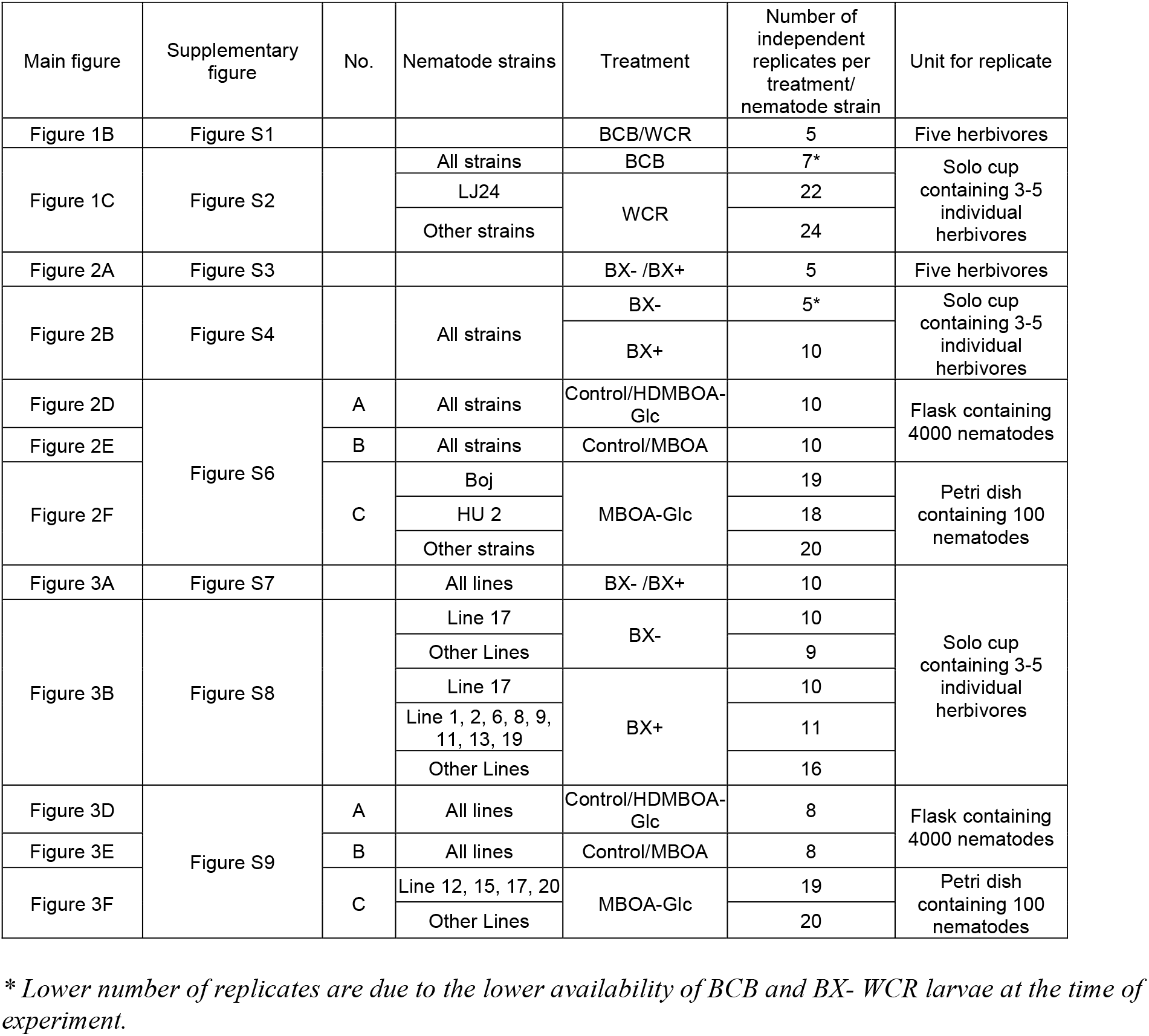
Numbers of individual biological replicates measured in the different experiments

| Main figure | Supplementary figure | No. | Nematode strains | Treatment | Number of independent replicates per treatment/ nematode strain | Unit for replicate |
| --- | --- | --- | --- | --- | --- | --- |
| Figure 1B | Figure S1 |  |  | BCB/WCR | 5 | Five herbivores |
| Figure 1C | Figure S2 |  | All strains | BCB | 7* | Solo cup containing 3-5 individual herbivores |
|  |  |  | LJ24 | WCR | 22 |  |
|  |  |  | Other strains |  | 24 |  |
| Figure 2A | Figure S3 |  |  | BX- /BX+ | 5 | Five herbivores |
| Figure 2B | Figure S4 |  | All strains | BX- | 5* | Solo cup containing 3-5 individual herbivores |
|  |  |  |  | BX+ | 10 |  |
| Figure 2D | Figure S6 | A | All strains | Control/HDMBOA-Glc | 10 | Flask containing 4000 nematodes |
| Figure 2E |  | B | All strains | Control/MBOA | 10 |  |
| Figure 2F |  | C | Boj | MBOA-Glc | 19 | Petri dish containing 100 nematodes |
|  |  |  | HU 2 |  | 18 |  |
|  |  |  | Other strains |  | 20 |  |
| Figure 3A | Figure S7 |  | All lines | BX- /BX+ | 10 | Solo cup containing 3-5 individual herbivores |
| Figure 3B | Figure S8 |  | Line 17 | BX- | 10 |  |
|  |  |  | Other Lines |  | 9 |  |
|  |  |  | Line 17 | BX+ | 10 |  |
|  |  |  | Line 1, 2, 6, 8, 9, 11, 13, 19 |  | 11 |  |
|  |  |  | Other Lines |  | 16 |  |
| Figure 3D | Figure S9 | A | All lines | Control/HDMBOA-Glc | 8 | Flask containing 4000 nematodes |
| Figure 3E |  | B | All lines | Control/MBOA | 8 |  |
| Figure 3F |  | C | Line 12, 15, 17, 20 | MBOA-Glc | 19 | Petri dish containing 100 nematodes |
|  |  |  | Other Lines |  | 20 |  |

\* Lower number of replicates are due to the lower availability of BCB and BX- WCR larvae at the time of experiment.

**Table S3.**
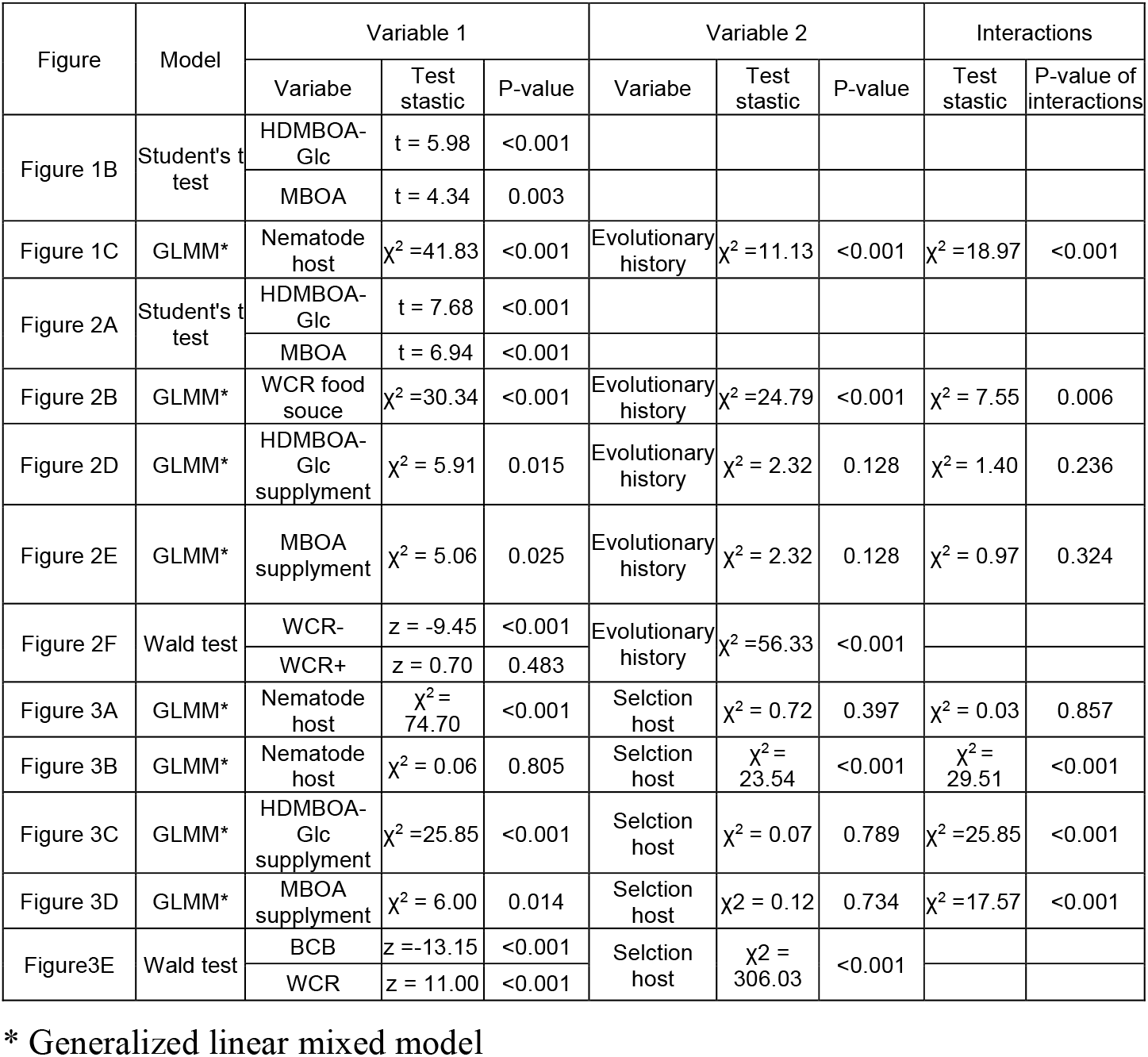
Summary of statistical models for main factors and interactions

**Table S4.** Summary of model selection procedures for multivariate analyses

| Intercept | HDMBOA-Glc toxicity (HG) | MBOA toxicity (M) | MBOA-Glc repellency (MG) | HG x M | HG x MG | M x MG | AICc | ΔAICc | w | R<sup>2</sup> |
| --- | --- | --- | --- | --- | --- | --- | --- | --- | --- | --- |
| 0.910 |  |  | -1.665 |  |  |  | -27.9 | 0.00 | 0.547 | 0.374 |
| 0.852 | 0.258 |  | -1.565 |  |  |  | -25.5 | 2.39 | 0.165 | 0.385 |
| 0.884 |  | 0.156 | -1.619 |  |  |  | -25.2 | 2.75 | 0.138 | 0.376 |
| 0.729 | 2.250 |  | -1.304 |  | -4.424 |  | -22.8 | 5.14 | 0.042 | 0.395 |
| 0.860 | 0.484 | -0.351 | -1.579 |  |  |  | -22.5 | 5.37 | 0.037 | 0.390 |
| 0.847 |  | 1.091 | -1.539 |  |  | -2.163 | -22.0 | 5.87 | 0.029 | 0.377 |
| 0.729 | 2.654 | -0.405 | -1.302 |  | -4.741 |  | -19.5 | 8.42 | 0.008 | 0.401 |
| 0.717 | 0.710 | 2.776 | -1.271 |  |  | -7.783 | -19.4 | 8.48 | 0.008 | 0.399 |
| 0.874 | 0.382 | -0.506 | -1.606 | 1.809 |  |  | -19.2 | 8.75 | 0.007 | 0.393 |
| 0.167 |  |  |  |  |  |  | -18.8 | 9.10 | 0.006 | 0.000 |
| 0.129 | 0.679 |  |  |  |  |  | -18.6 | 9.35 | 0.005 | 0.090 |
| 0.139 |  | 0.693 |  |  |  |  | -17.7 | 10.23 | 0.003 | 0.057 |
| 0.130 | 0.761 | -0.125 |  |  |  |  | -15.7 | 12.20 | 0.001 | 0.090 |
| 0.740 | 2.507 | -0.440 | -1.324 | 0.443 | -4.474 |  | -15.6 | 12.33 | 0.001 | 0.401 |
| 0.735 | 3.018 | -1.092 | -1.315 |  | -5.644 | 1.686 | -15.6 | 12.34 | 0.001 | 0.401 |
| 0.727 | 0.678 | 2.576 | -1.293 | 0.345 |  | -7.359 | -15.5 | 12.40 | 0.001 | 0.399 |
| 0.129 | 0.814 | -0.042 |  | -0.986 |  |  | -12.6 | 15.33 | 0.000 | 0.091 |
| 0.754 | 3.094 | -1.638 | -1.354 | 0.570 | -5.958 | 2.915 | -11.2 | 16.74 | 0.000 | 0.401 |

Model selection for Fig. 3
| Intercept | HDMBOA-Glc toxicity (HG) | MBOA Toxicity (M) | MBOA-Glc repellency (MG) | HG x M | HG x MG | M x MG | AICc | ΔAICc | w | R<sup>2</sup> |
| --- | --- | --- | --- | --- | --- | --- | --- | --- | --- | --- |
| 0.505 |  |  | -0.983 |  |  |  | -27.0 | 0.00 | 0.423 | 0.650 |
| 0.378 | 0.578 |  | -0.792 |  |  |  | -26.0 | 1.05 | 0.251 | 0.685 |
| 0.516 |  | -0.083 | -1.002 |  |  |  | -23.9 | 3.16 | 0.087 | 0.650 |
| 0.434 | -0.918 |  | -0.892 |  | 3.667 |  | -23.6 | 3.43 | 0.076 | 0.704 |
| 0.440 | 0.731 | -0.697 | -0.899 |  |  |  | -23.1 | 3.96 | 0.058 | 0.696 |
| 0.479 |  | 3.058 | -0.969 |  |  | -7.371 | -21.7 | 5.32 | 0.030 | 0.675 |
| 0.515 | -0.922 | -0.838 | -1.033 |  | 4.129 |  | -20.5 | 6.53 | 0.016 | 0.719 |
| 0.531 | -2.548 | 4.756 | -1.114 |  | 7.893 | -13.190 | -20.2 | 6.79 | 0.014 | 0.777 |
| 0.414 | 0.675 | 1.995 | -0.879 |  |  | -6.204 | -20.0 | 6.98 | 0.013 | 0.713 |
| 0.219 | 2.175 | 9.359 | -0.602 | -29.360 |  | -19.380 | -19.9 | 7.10 | 0.012 | 0.774 |
| 0.414 | 1.018 | -0.405 | -0.857 | -5.205 |  |  | -19.2 | 7.85 | 0.008 | 0.700 |
| -0.072 | 1.562 |  |  |  |  |  | -17.5 | 9.50 | 0.004 | 0.437 |
| 0.781 | -5.418 | -2.536 | -1.489 | 25.560 | 11.840 |  | -17.3 | 9.72 | 0.003 | 0.742 |
| -0.073 | 1.144 | 1.017 |  |  |  |  | -15.6 | 11.46 | 0.001 | 0.470 |
| 0.408 | -0.774 | 6.817 | -0.916 | -12.150 | 5.070 | -16.150 | -14.7 | 12.35 | 0.001 | 0.779 |
| -0.043 |  | 2.372 |  |  |  |  | -14.6 | 12.38 | 0.001 | 0.350 |
| -0.077 | 1.969 | 1.670 |  | -16.010 |  |  | -13.7 | 13.28 | 0.001 | 0.516 |
| 0.012 |  |  |  |  |  |  | -8.8 | 18.19 | 0.000 | 0.000 |

Summary of the averaged model:
| Term | Estimate | SE | Relative importance |
| --- | --- | --- | --- |
| Intercept | 0.458 | 0.121 | - |
| MBOA-Glc repellency | -0.912 | 0.210 | 1.00 |
| HDMBOA-Glc toxicity | 0.215 | 0.379 | 0.37 |

*R<sup>2</sup> of the averaged model: 0.671*

